# Live Cell Fluorescence Microscopy – An End-to-End Workflow for High-Throughput Image and Data Analysis

**DOI:** 10.1101/2024.03.28.587214

**Authors:** Jakub Zahumensky, Jan Malinsky

## Abstract

Fluorescence microscopy images of biological samples contain valuable information but require rigorous analysis for accurate and reliable determination of changes in protein localization, fluorescence intensity and morphology of the studied objects. Traditionally, cells for microscopy are immobilized using chemicals, which can introduce stress. Analysis often focuses only on colocalization and involves manual segmentation and measurement, which are time-consuming and can introduce bias. Our new workflow addresses these issues by gently immobilizing cells using a small agarose block on a microscope cover glass. This approach is suitable for cell-walled cells (yeast, fungi, plants, bacteria), facilitates their live imaging under conditions close to their natural environment and enables the addition of chemicals during time-lapse experiments. The primary focus of the protocol is on the presented analysis workflow, which is applicable to virtually any cell type – we describe cell segmentation using the Cellpose software followed by automated analysis of a multitude of parameters using custom-written Fiji (ImageJ) macros. The results can be easily processed using the provided R markdown scripts or available graphing software. Our method facilitates unbiased batch analysis of large datasets, improving the efficiency and accuracy of fluorescence microscopy research.

The reported sample preparation protocol and Fiji macros were used in our recent publications: *Microbiol Spectr* (2022), DOI: 10.1128/spectrum.01961-22; *Microbiol Spectr* (2022), DOI: 10.1128/spectrum.02489-22; *J Cell Sci* (2023), DOI: 10.1242/jcs.260554.

**Graphical overview:** 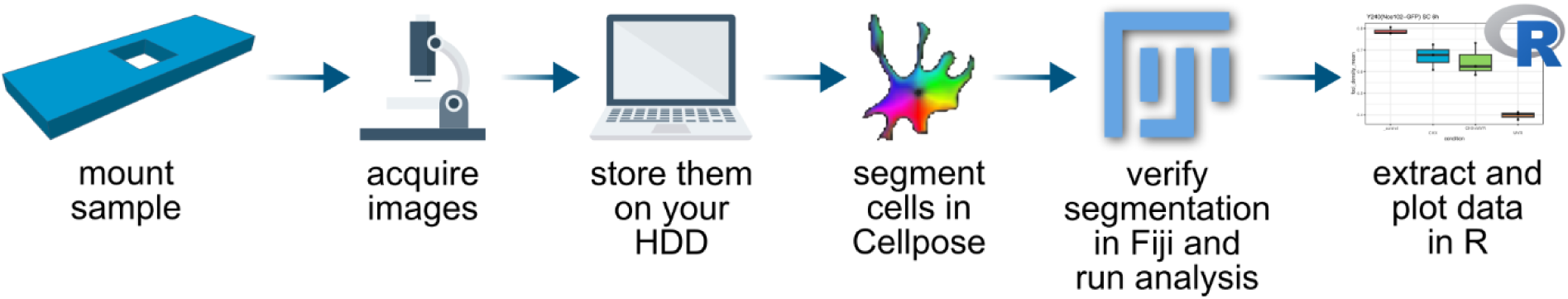

***From fluorescence microscopy to numbers and plots – a generalized workflow***

## Introduction

Well-acquired microscopy images of biological samples contain high amounts of valuable information. However, the space available in scientific papers is generally insufficient to display multiple images for each biological replicate. Therefore, only a single representative image, often of a single cell, is usually shown. This results in a lot of important information not being communicated to the reader. This can be redeemed by performing a formal rigorous statistical analysis of all acquired data and reporting information on variation across biological replicates and cells in the form of plots and statistical analysis. In this way, it is readily clear how many times an experiment was performed, how many cells were analyzed in each replicate, and how significant the reported differences between strains/cell lines and/or conditions are. When the formal analysis of microscopy images is performed, it is often limited to the quantification of colocalization where applicable. When additional analysis is performed, the cells are segmented and analyzed manually (e.g.,^1^). However, any user-dependent steps limit the analysis and can introduce bias and errors.

For the preparation of samples for live cell microscopy, concanavalin A or poly-L-lysine is often used to help cells adhere to glass surfaces via physical interactions. In our approach, (cell-walled) cells in thick suspensions are placed on a microscopy cover glass and overlaid with a small block of 1% agarose prepared either in buffer or cultivation medium identical to that of the respective cell culture, ensuring a close-to-natural environment. The cover glass is taped to a custom-made microscopy sample holder (Fig. 1A, 2). In this approach, the cells are gently immobilized on the cover glass just by mild mechanical forces, which does not induce a cellular stress response, as documented by the normal actin structure in *C. elegans*^2^ and by the absence of stress granules *S. cerevisiae* cells^3,4^. Furthermore, the cells spontaneously prefer a monolayer arrangement, i.e., they are mostly in the same focal plane. This makes microscopy imaging more efficient, saving both time and storage space. Since agarose is permeable to small molecules, cells can be treated by directly adding chemicals to the agarose block (when using an inverted microscope) during time-lapse experiments^3–5^. Adding a cover glass on the other side of the sample holder (Fig. 2) prevents water evaporation and shrinkage of the agarose block, enabling long time-lapse experiments. The described arrangement thus represents an alternative to both surfaces coated with concanavalin A or poly-L-lysine and expensive microfluidics systems.

**Figure 1.**
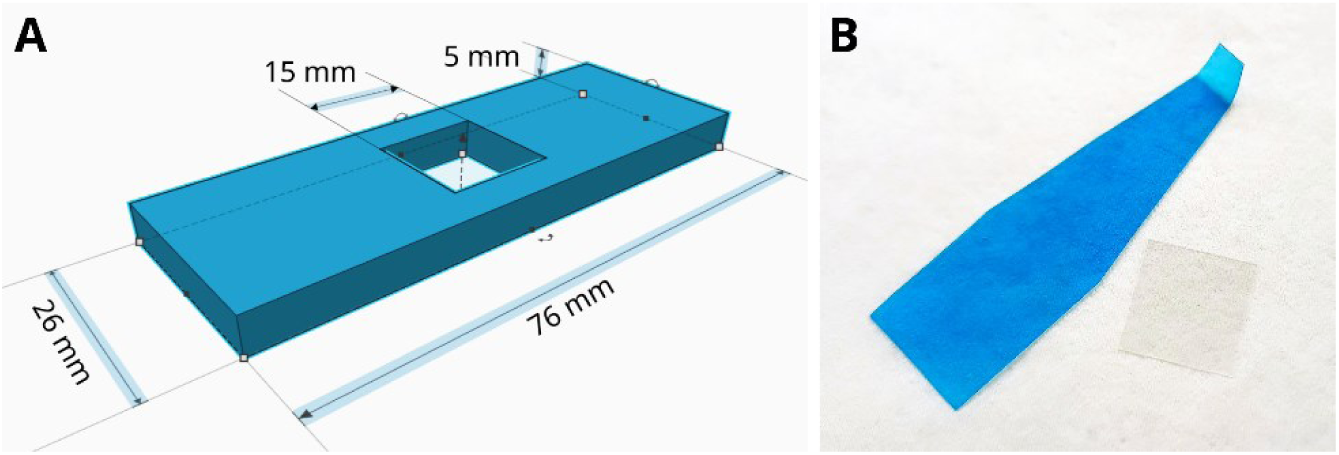
Equipment for microscopy sample preparation. A) Image of the custom-made microscopy sample holder; the Tinkercad file for 3D printing can be downloaded here: https://www.tinkercad.com/things/0gmLJyrrJ3z?sharecode=lbC9EIcwjEp6yeSq8FMaTI7W8Y652L81cd3yImyDp1k. B) A spatula made of old Western blot film (a 22×22 mm cover glass for scale).

**Figure 2.**
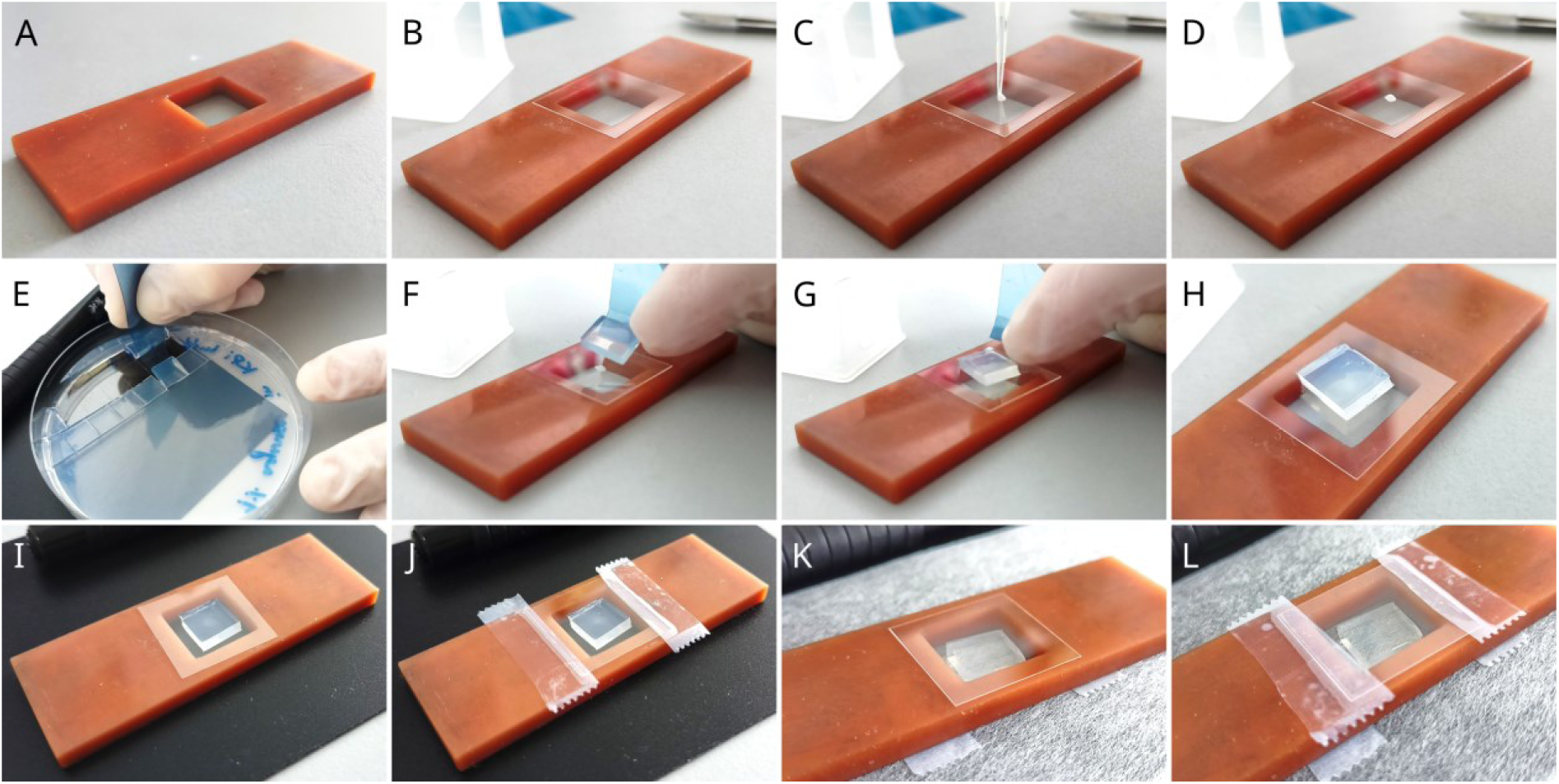
Microscopy specimen preparation. A) The sample holder. B) Place cover glass on a level surface, e.g., the sample holder. C-D) Pipette 1 μL of the cell suspension on the cover glass. E) Use the spatula to lift a small block (∼1×1 cm) of pre-cut agarose from the Petri dish. F-H) Cover the cell suspensions with the agarose block. I) Turn the glass with agarose upside down so that the agarose block is inside the square hole in the sample holder. J) Tape the cover glass to the holder. K-L) For time-lapse experiments, turn the sample holder upside down (putting it on a non-abrasive surface) and tape another cover glass to seal the chamber.

The analysis protocol reported here integrates powerful automatic segmentation using Cellpose software^6^ with automatic cell-by-cell analysis performed by custom Fiji (ImageJ) macros. This approach allows for the processing of large microscopy datasets from unrelated experiments in batch mode, in a consistent and unbiased fashion. In addition to the initial verification of proper segmentation (which can be blinded), the analysis does not require user input. It can, therefore, be run unassisted. Data from individual experiments can then be easily extracted and analyzed to generate graphs and perform statistical analyses using provided custom R scripts. Alternatively, the Results table can be processed manually using the Multiple variables data table in the GraphPad Prism graphing software, or in Microsoft Excel/LibreOffice Calc – or by writing one’s own macros. We have successfully used the reported protocol for sample preparation and Fiji macros in our recent publications: *Microbiol Spectr* (2022), DOI: 10.1128/spectrum.01961-22^7^; *Microbiol Spectr* (2022), DOI: 10.1128/spectrum.02489-22^8^; *J Cell Sci* (2023), DOI: 10.1242/jcs.260554^9^.

## Materials and Methods

### Biological materials

The microscopy sample preparation protocol described below was developed for live microscopy of *S. cerevisiae* and *C. albicans* cells (verified suppliers of yeast are Euroscarf or Dharmacon; choose depending on your location), but its use is not limited to these two yeast species. It is applicable for all cell types that have a cell wall, as we have previously tested with the cells of the *Ensifer meliloti* bacterium and *Nicotiana tabacum* plant. The subsequent microscopy data analysis works with images of all cell types irrespective of organism and/or tissue, and labeling.

### Agarose blocks for microscopy sample preparation

In our experimental setup we use 1% agarose prepared either using a potassium phosphate (KP_i_) buffer or cultivation medium (see below) to gently immobilize cell-walled cells on the microscopy cover slips. We have previously tested various concentrations of agarose and found 1% to be the most suitable for this purpose. Agarose of lower concentrations is brittle and hard to manipulate. The use of higher agarose concentrations, on the other hand, results in a refractive index mismatch on the agarose block/medium interface, which causes undesirable light reflections during imaging. In addition, denser agarose is inflexible, which again leads to difficult manipulation.

For general use, 1% nonfluorescent agarose in 50 mM potassium phosphate (KPi) buffer (pH 6.3) is prepared by mixing 7.75 ml of 1 M KH_2_PO_4_ with 2.25 ml of 1 M K_2_HPO_4_ and mixing it with distilled water to 200 ml. After boiling, 10 ml of this solution was poured into a 90 mm diameter Petri dish (and stored at 4°C for up to a month after the petri dish was sealed with Parafilm). The agarose was cut into ∼1×1 cm squares with a scalpel blade, and a small spatula (Fig. 1B) was used to manipulate the blocks.

For specific purposes (e.g., time-lapse experiments), 1% conditioned agarose was made as follows: When harvesting (yeast) culture by centrifugation, 3 mL of the supernatant was removed into a fresh 15 mL Falcon tube containing 0.03 g of agarose. The mixture was closed firmly, placed in a tall glass beaker and heated carefully at small intervals in a microwave oven at a low intensity until the agarose had dissolved completely. After cooling down to touch, chemicals were added as required for the respective experiment, and the mixture was poured into a 35 mm diameter Petri dish placed on a level surface. Using conditioned agarose ensured that the environment in which the cells were grown was retained, minimizing stress exposure. This is especially important when changes in either the composition of the media or its pH can result in significant changes in the localization, shape, or amount of the structure of interest and when the budding of yeast cells is to be monitored^1,3,7^.

### Microscopy sample preparation

The following procedure uses budding yeast as an example but is suitable for any cells with a cell wall. The centrifugation settings need to be adjusted based on the cell size of the used organism.

1. Cultivate cells in liquid media under desired conditions.
2. Transfer the whole culture to a 50 ml Falcon tube.
3. Centrifuge for 2 min at 1430 rcf at room temperature.
4. Remove most of the supernatant by pipetting or decanting.
5. Resuspend the pellet using a vortex mixer in 3 one-second pulses (850 rpm)
Alternatively, keep the supernatant and disturb the pellet gently by pipetting – especially useful when low-density cultures (for example, exponentially growing yeast) are processed.
6. Pipette 1 μL of the cell suspension onto a clean 0.17 mm thick microscopy cover glass.
7. Cover the cell suspension with a block of 1% agarose block prepared with either KP_i_ buffer or cultivation media (conditioned agarose, see above), using the small flat spatula (Fig. 1B) to manipulate the agarose block.
8. Place the cover glass into the sample holder (Fig. 1A; a Tinkercad file for 3D printing can be downloaded here: https://www.tinkercad.com/things/0gmLJyrrJ3z?sharecode=lbC9EIcwjEp6yeSq8FMaTI7W8 <u>Y652L81cd3yImyDp1k</u>), with the agarose block inside the chamber.
9. Tape the cover glass to the holder with Scotch tape.
10. *Optional:* For time-lapse experiments and when a microscope stage incubator is used to control temperature, flip the sample holder upside down onto a lens cleaning paper and tape another cover glass onto the other side (this prevents the agarose block from drying out during longer experiments).

*Notes: Steps 6-10 are depicted in Fig. 2. A standalone version of the sample preparation protocol (including material) is available at protocols.io:* https://www.protocols.io/view/live-cell-microscopy-sample-preparation-yeast-cult-8epv5r23dg1b^10^.

While this procedure greatly facilitates the spontaneous organization of cells into a monolayer, areas with gaps and/or multilayer arrangement will likely also be present in the sample. We provide examples of these in Fig. 3 and recommendations on tackling issues in the Troubleshooting section of the Discussion.

**Figure 3.**
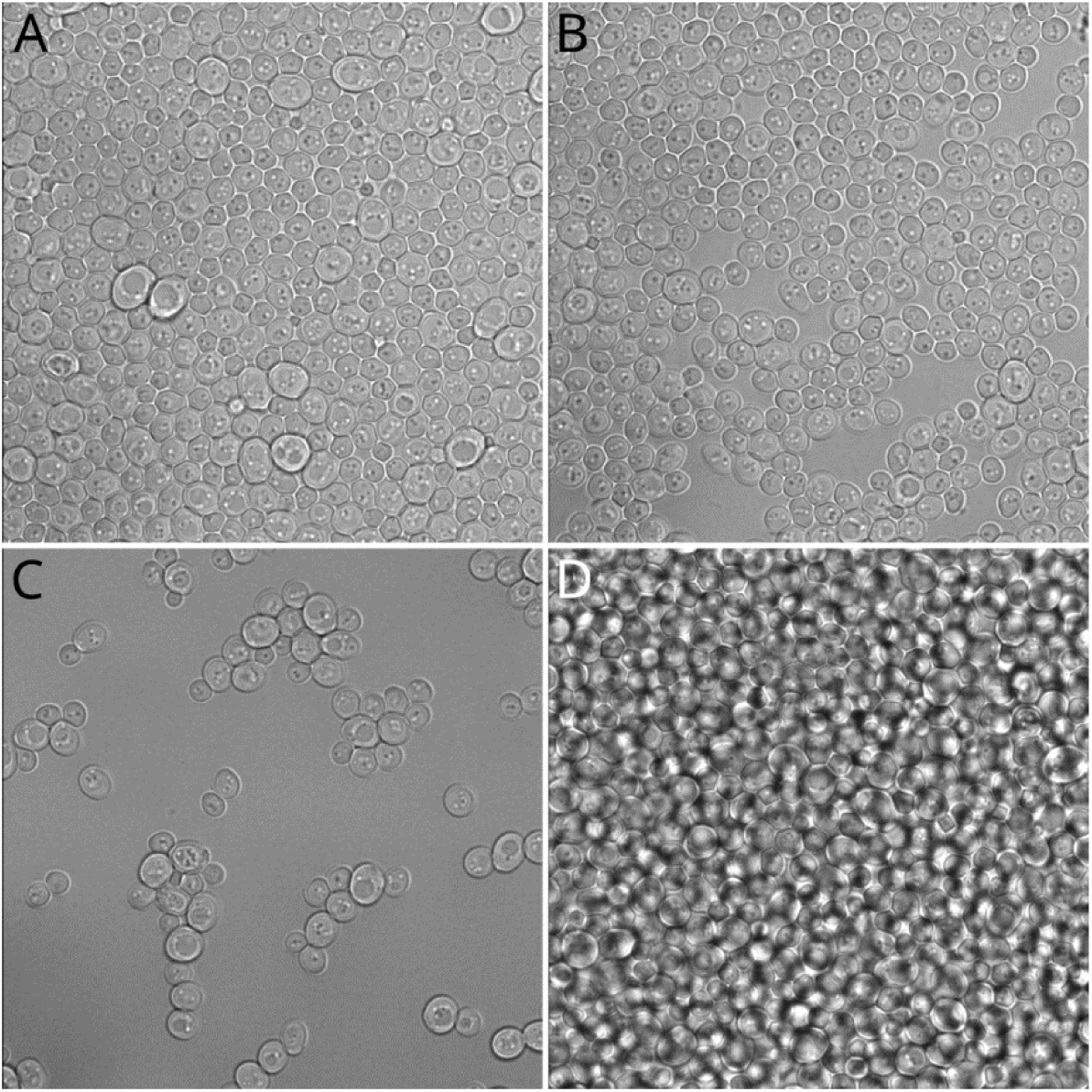
Visual guide for microscopy sample preparation success. Depending on the turbidity of the cell suspension, four different types of cellular arrangement on the cover slip emerge spontaneously: A) Monolayer – ideal for imaging and subsequent analysis. B) Monolayer with gaps – not ideal, but workable. C) Widely spaced groups of cells and individual cells – should be avoided, unless cell division is to be monitored over the course of multiple generations. D) Multilayer – undesirable. For optimization refer to the Troubleshooting section.

### Microscopy image acquisition for analysis

Setting up the microscope for imaging usually requires certain compromises and trade-offs. While using a higher laser power generally leads to a better signal-to-noise ratio, it can cause phototoxicity and sample heating, which is not compatible with time-lapse experiments. Imaging with high spatial resolution limits the rate at which time series can be acquired. When higher temporal resolution is required, signal and/or spatial resolution has to be sacrificed. For longer experiments, the intensity of the excitation light typically needs to be decreased to avoid photobleaching (Fig. 4). Understanding these aspects is important and the experimenter needs to carefully consider each aspect to decide which is the most crucial for their experiment.

**Figure 4.**
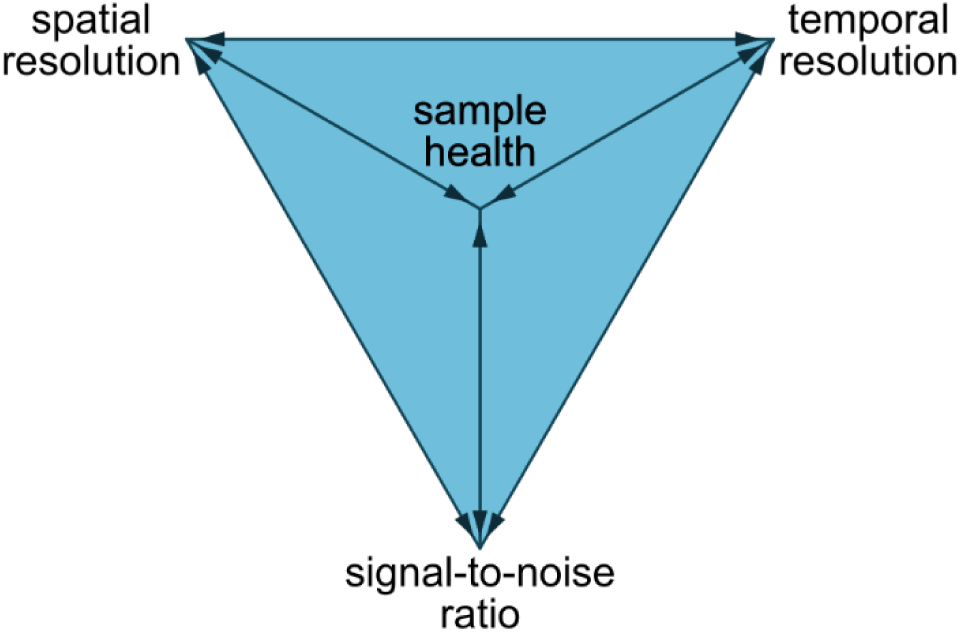
The trade-off triangle. When setting up a microscope for a specific experiment, a certain amount of compromise is required due to physical limitations of both the system and the sample. See text for details.

Regardless of this, there are certain general rules that should be followed when acquiring microscopy images. This is especially true when downstream analysis is to be applied. Failure to adhere to these can render the analysis results worthless or simply wrong (for a more detailed discourse see e.g.,^11^):

1. Avoid overexposure of images.
*Note: Detector saturation leads to incorrect intensity readings and obscuring of bright features of the image. In some instances, this is inevitable (e.g., when axons and dendrites are to be imaged in a neuron with high-intensity body and one is not concerned with the body), but such images are not suitable for the analysis presented in this protocol.*
2. Set offset properly (when using a photomultiplier tube as the detector).
*Note: The images need to be acquired in a way that the background intensity is low, but not zero. Only a few, and clearly separated, zero-intensity pixels should be present in a single image. Failure to adhere to this will lead to incorrect intensity readings from the image.*
3. Take care not to bleach the fluorescent tags.
*Note: Violations of any of 1-3 result in incorrect intensity readings*
*Note: You can minimize bleaching by finding the focus in one image area and imaging another area directly adjacent, e.g., using the tile scan option of your microscope or moving the stage in a defined manner using the stage controller in the microscope software.*
4. Strive to minimize noise.
*Note: Microscopy imaging always requires that certain compromises be made (Fig. 4) and noise is always present in microscopy images. However, good signal-to-noise ratios are highly desirable for downstream analysis, as high noise levels can obscure subcellular structures and cause issues during the segmentation of cells.*
5. Avoid bleed-through (crosstalk).
*Note: Bleed-through is caused by an overlap of emission spectra of two or more fluorescent markers. It can be easily mistaken for colocalization and hence lead to misinterpretation of the data. Using sequential acquisition of the signal from individual channels greatly reduces the risk of bleed-through*.
6. Do not change acquisition settings between imaging samples of the same series.
*Note: Changing the settings results in different intensities caused by the instrumentation, not the sample (strain/cell type, condition). As a result, a comparison of fluorescence intensities among the samples is impossible. Note that doubling the laser intensity generally does not result in doubling the fluorescence intensity.*
7. Do not directly compare intensities from samples obtained on different days.
*Note: Like 6, but more easily overlooked. In general, it is better to use intensity ratios of values originating from a single sample (e.g., ratio of the mean intensity in the plasma membrane to that in the cytosol), as these are based on an internal reference. Absolute fluorescence intensity readouts across samples should be compared only as a guide, and only from samples imaged in an immediate succession. Intensity readings are unreliable due to day-to-day fluctuations in the current, temperature, laser intensity, etc. In addition, to compare intensities across replicates, the intensities need to be first normalized – see Discussion.*

## Data analysis

Here, we report our custom Fiji (ImageJ) macros that facilitate automatic analysis of large numbers of segmented objects, such as cells, reporting a multitude of parameters for each, including size (area), fluorescence intensity (both integral and mean) and the number, fluorescence intensity and distribution of microdomains (areas with high local fluorescence intensity, also referred to as high-intensity foci) in the border areas of the objects. Shape descriptors are included (major, minor ellipse axis and eccentricity). The fluorescence intensity of the objects is reported separately for the whole objects, their circumference and for their inside. When whole cells are analyzed, this translates to intensity readings from the whole cell, the plasma membrane, and the intracellular space. In addition, various ratios are calculated and reported. The macros process all images of all biological replicates of all experiments found within the specified folder and all its subfolders (i.e., they work recursively). For the down-stream processing of results to work properly, data from various biological replicates need to be stored in separate folders and named consistently (see below). The data are reported in a single Results table where each line contains information on a single cell (analyzed object), including which experiment and biological replicate it belongs to.

While the microscopy sample preparation using agarose blocks described above is useful only for microscopy of cell-walled cells (yeast and other fungi, plants and bacteria), the data analysis macros can be used without issues with any type of microscopy images – live or fixed cells, yeast, mammalian cultures, etc. As long as the images are acquired in accordance with the guidelines for microscopy imaging described above and the segmentation (see below) is performed properly, the analysis will return sensible data.

The analysis workflow itself has the following steps, which are described in detail below:

1. Software preparation and data organization

1.1. Install required software, Fiji plugins and macros (Fig. 5)
1.2. Store acquired images in a proper file structure, according to Fig. 6.
2. Preparation for analysis – image export and object segmentation

2.1. In Fiji, load and run the “1. export.ijm” macro to export the images to tiff format.
2.2. In Cellpose, open one of the exported images and find segmentation parameters; close Cellpose.
2.3. In the Anaconda command line, run the Cellpose batch segmentation command (see below for an example) updated with required segmentation parameters.
3. Preparation for analysis – regions of interest (ROI) preparation and segmentation verification

3.1. In Fiji, load and run the “2. ROI prep.ijm” macro and select the “Convert Masks to ROIs” option to convert the Cellpose-generated segmentation masks to ROIs.
3.2. Run the “2. ROI prep.ijm” macro again and select “Check ROIs” option. Audit the images one by one to verify that the segmentation is correct and adjust the ROIs as necessary.
4. Running the analysis

4.1. In Fiji, load and run the “3. quantify.ijm” macro to perform the actual analysis.
4.2. View the Results table.
5. Extracting numbers, plotting and performing statistical analysis

5.1. In R Studio, load the provided R markdown. Run the R scripts using the “Run” button at the top, or by pressing “ctrl+alt+R”.
Alternatives: use GraphPad Prism, Systat Sigmaplot, Microsoft Excel/LibreOffice Calc

**Figure 5.**
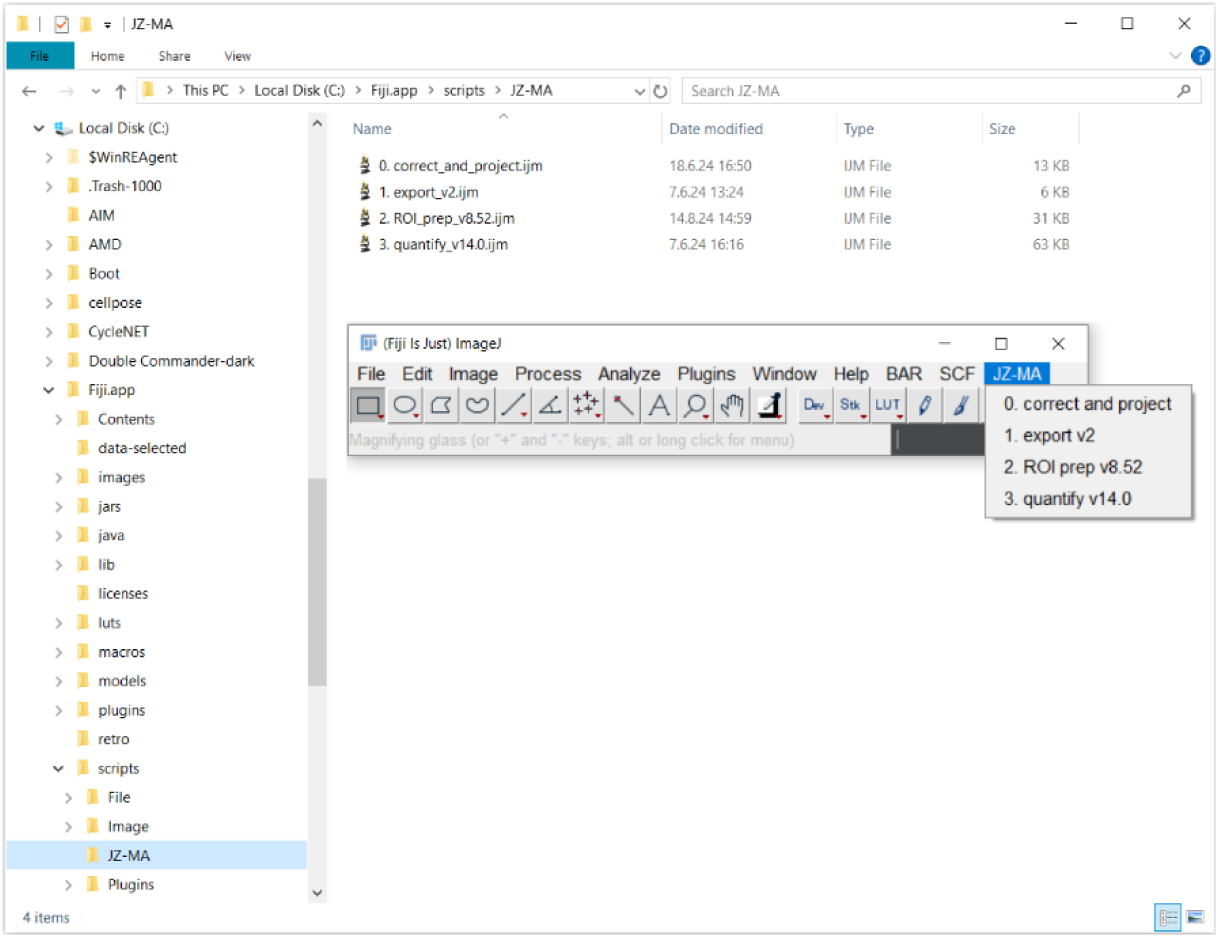
Making the macros available from a Fiji menu. After placing the macros into the indicated subfolder in the “Fiji.app/scripts/” folder, they can be run easily from the newly created menu in the main Fiji window. Note that Fiji needs to be restarted for the changes to take effect.

**Figure 6.**
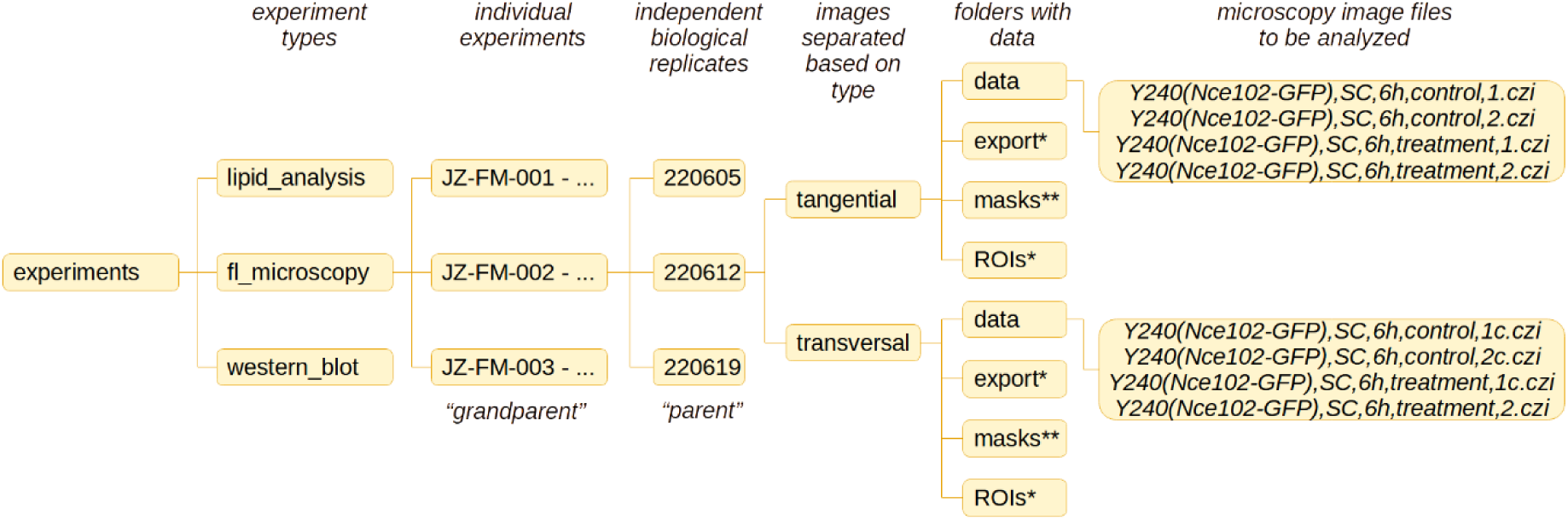
Example data organization structure. The folders marked with a single and double asterisk are created automatically by the provided Fiji macros and Cellpose, respectively.

### Step 1: Software preparation and data organization

#### Step 1.1: Installation of required software and the use of macros

The following macros and software are either required or recommended (marked as optional) for the microscopy image analysis described below:

- Our custom Fiji macros and R scripts written for this protocol; deposited to GitHub: https://github.com/jakubzahumensky/microscopy_analysis<u>; their use is described below</u>
- Cellpose 2.0 (Stringer et al., 2021): https://www.cellpose.org/; installation instructions at https://github.com/MouseLand/cellpose/blob/main/README.md

The Cellpose Readme website includes instructions on the installation of Python (https://www.python.org/downloads/) and Anaconda (https://www.anaconda.com/download); both are required for the running of Cellpose
- Fiji (ImageJ 1.53t or higher; https://imagej.net/software/fiji/downloads) with additional plugins/libraries that are not part of the general Fiji download:

- SCF-MPI-CBG and BIG-EPFL (to install, navigate through Help → Update → Manage update sites, then tick checkboxes for BIG-EPFL and SCF MPI CBG. Click Apply and Close → Apply changes → OK; restart Fiji)
- Watershed_ plugin^12^ (to install, download from http://bigwww.epfl.ch/sage/soft/watershed/ and put in the plugins folder of Fiji/ImageJ)
- Adjustable Watershed plugin (to install, download Adjustable_Watershed.class from https://github.com/imagej/imagej.github.io/blob/main/media/adjustable-watershed and put it in the Plugins folder of Fiji/ImageJ)
- Up-to-date version of R (including R Studio) – recommended for automatic data processing
- Prism 8 or higher (GraphPad) – *optional* – for semi-automatic data processing
- Microsoft Excel/LibreOffice Calc – *optional* – for manual data processing (not recommended)

The simplest way to make the macros available in Fiji and to use them afterwards is to create a menu in the main Fiji application window. This is done by creating a subfolder in the “Fiji.app/scripts/” folder of the user’s local Fiji installation and placing the macro files into this subfolder (note that the filenames must not contain spaces). The name of this subfolder can be chosen by the user and will be displayed as the title of the menu with macros. Here, we use the folder name “JZ-MA” as an example (Fig. 5).

Alternatively, all reported macros can be run from Fiji by clicking “Plugins” → “Macros” → “Run…” and opening the macro from the shown dialog window or by drag-and-dropping the respective macro onto the Fiji window and pressing “Run” in the macro edit window. The macros are numbered in the order in which they are to be used. Each macro run starts by displaying a dialog window, where multiple parameters need to be set by the user. The path to the folder to be processed can be copy-pasted or selected using the “Browse” button. All dialog windows displayed by the macros include a “Help” button, which shows information on individual parameters. The user is informed when the macro run is finished with a message (“Finito!”). If there are any irregularities during ther macro run, the message also provides a warning and additional information (e.g., that not all images have defined regions of interest, with their list written in one of the log windows).

#### Step 1.2: Required data organization

For the analysis to work, a certain data organization structure is required (Fig. 6). If you plan to process data from multiple experiments at the same time, it is advantageous to use experimental (accession) codes for your experiments. These codes also facilitate cross-referencing between digital data and hard-copy lab notebooks (as well as between related experiments). We use experiment codes following the “JZ-FM-αβγ” format, where JZ are initials of the experimenter (here, “Jakub Zahumensky”, FM denotes “fluorescence microscopy” and αβγ is a three-digit number). All the data belonging to a specific experiment should be stored in a single folder whose name starts with the experiment code. Within this folder, each biological replicate should have a separate folder where data are stored. We recommend using the date in the YYMMDD format to keep the folders ordered chronologically. In each replicate folder, separate folders for transversal (images of cell cross-sections) and tangential (depicting cell surfaces) images need to be created, as required. In these, a folder called “data” is used for storing the actual data to be analyzed. All additional folders that are required are created by the provided macros. If images require preprocessing, such as drift correction or z-projection, the use of an additional “data-raw” (or similar) folder is recommended. The actual filenames should include the following information: cell type (yeast strain/cell line etc.), fluorescent tag(s) listed in the order in which channels are saved in multichannel images, cultivation medium and temperature (if multiple options exist) and time, condition/treatment and frame number (multiple frames of each strain and condition are required). Information on what part of cells is imaged (transversal/tangential section, z-stack) should be included as well. Images from time-lapse experiments and z-stacks need to be corrected for drift and bleaching (e.g., using the “0. correct and project.ijm” macro) before the analysis. Time-series also need to be separated into individual images. As the Results table is comma-separated, the information in filenames also needs to be comma-separated, ideally without spaces (example in Fig. 6; any spaces are substituted with underscores in the Results table). The experiment codes, biological replicate folder names and filenames need to be used consistently (including using the exact same notation for the same treatment each time), as these are reported in the Results table and enable automatic data filtering and processing (using R/GraphPad Prism), as well as manual processing (Microsoft Excel/LibreOffice Calc). Experiments destined for analysis can be temporarily moved to a single folder and analyzed in a single run.

### Step 2: Preparation for analysis – image export and object segmentation

To segment objects (typically cells and/or nuclei) in microscopy images, we use Cellpose 2.0^6^ (see Software). In our experience, its use is reasonably simple, and we consistently obtain satisfactory segmentation results. Other segmentation options are available in Fiji (ImageJ) or Cell Profiler^13^, but these options are not covered in this protocol. Cellpose 2.0 can be used either through its graphical user interface (GUI) or via the command line (Anaconda; see Software). The GUI is useful for determining the segmentation parameters for a given cell type and experiment. The command line can then be used to run the segmentation in batch mode.

#### Step 2.1: Batch export of images using a Fiji macro

To prepare the images for segmentation, they need to be first exported into “tif” format, as Cellpose does not work with raw microscopy images. To export all images across biological replicates (and/or experiments), run the “1. export.ijm” macro. For export, there is an option to adjust the contrast of the images, which might be beneficial for the following segmentation in Cellpose (note that if the contrast is adjusted, the images should be used only for segmentation, not for the analysis itself). The macro allows the user to specify which channels are to be exported. The segmentation masks and, in turn, ROIs are then used to quantify both channels. Therefore, it is advisable to export channels with brighter fluorescence signals and clearly visible plasma membranes. The analysis (described below) processes a single frame at a time. Therefore, for time-lapse experiments, images need to be split into individual frames.

#### Step 2.2: Determination of segmentation parameters using Cellpose GUI

To perform segmentation, follow instructions on the official Cellpose Readme website ((https://github.com/MouseLand/cellpose/blob/main/README.md; section “Run Cellpose locally”; includes a demo) to open the GUI. Open one of your exported images, and input the estimate of the diameter of your cells in the Segmentation tab. In the model zoo tab, select one of the models (cyto2 works well for yeast cells), which starts the segmentation. Once finished, change the flow_threshold (maximum allowed error of the flows for each mask), cellprob_threshold (higher values lead to smaller segmentation masks) and stitch_threshold (for segmentation of z-stacks) to find the parameters that give the best results (a detailed description of the parameters can be found in the Cellpose documenta-tion<u>-</u> https://cellpose.readthedocs.io/en/latest/settings.html). For the analysis macro (below) to work properly, the size of the Masks/ROIs for transversal images should be set in such a way that their edges are placed in the middle of the plasma membrane. For tangential images, masks should be just as large as the visible parts of the cells (Fig. 7). Once you have obtained the segmentation parameters (write them down), close the GUI.

**Figure 7.**
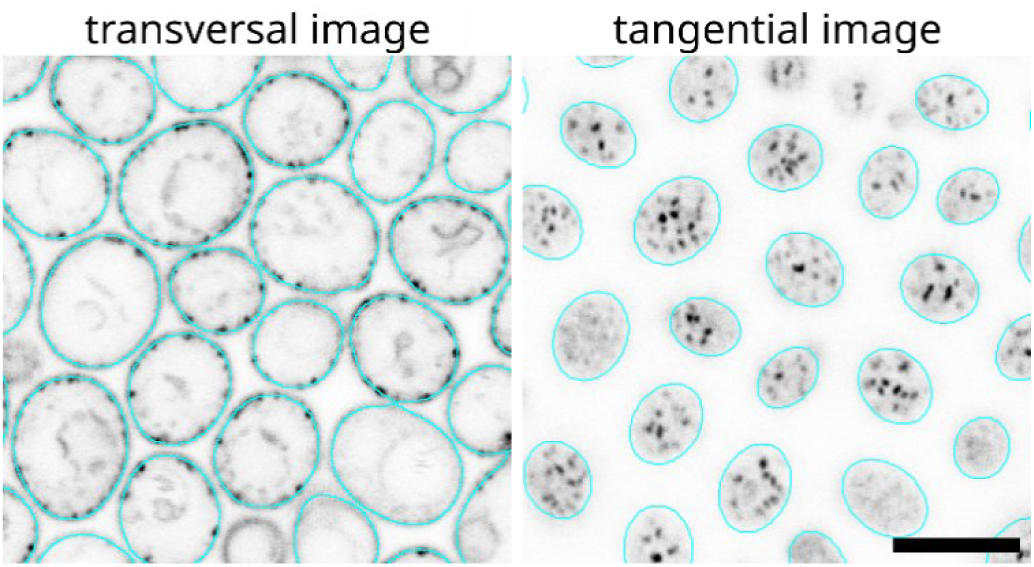
Guide to Segmentation. Segmentation should be performed in a way that the ROI edges are in the middle of the plasma membrane in transversal images (cell cross-sections) and just touch the outside of the objects in tangential images (cell surfaces). Yeast cultures were grown in synthetic com-plete media for 24 h and Nce102-GFP (a yeast plasma membrane microdomain protein) was imaged. Scale bar: 5 µm. Raw data from Zahumensky et al., 2022^7^.

#### Step 2.3: Batch segmentation using the Anaconda command line

To segment images in batch mode using Cellpose, use a variation of the following command in the Anaconda prompt (detailed information can be found in the Cellpose documentation -https://cellpose.readthedocs.io/en/latest/cli.html#cellpose-cli):

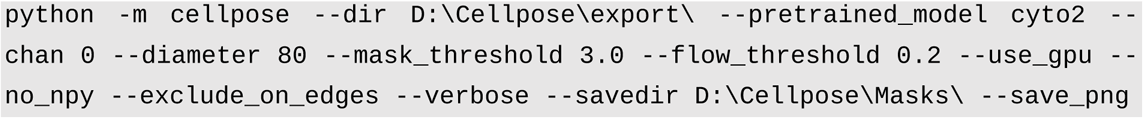

In the command, substitute the “D:\Cellpose” folder for where your exported images are. Additionally, substitute values for diameter, mask_threshold and flow_threshold based on the parameters you obtained using the GUI. Note that the export of masks does not work if there are any spaces in the path to your files. To circumvent possible issues, we find it best to have a dedicated folder for Cellpose segmentation (such as in the example), move the exported images there, and move them back to their original location after segmentation, together with the masks folder (needs to be called “masks” for downstream automatic processing). For the “--use_gpu” option to work (i.e., use the graphical processing unit for calculations; optional), additional steps are required before running the segmentation (see Cellpose documentation - https://cellpose.readthedocs.io/en/latest/installation.html#amd-gpu-rocm-installation).

#### Step 3: Preparation for analysis – regions of interest (ROIs) and segmentation verification

Before analysis, the segmentation mask images obtained in Cellpose need to be converted to regions of interest (ROIs) that Fiji can use, and the accuracy of these ROIs needs to be verified. To perform this, run the “2. ROI_prep.ijm” macro in Fiji. This macro is used for both tasks and needs to be run twice.

#### Step 3.1: Conversion of masks to ROIs using a Fiji macro

In the first macro run, select the “Convert Masks to ROIs” in the process option. This macro converts all Masks of the selected image type into ROIs and saves these in a newly created “ROIs” folder in the form of zip files called RoiSets (see Fig. 6). The process runs in batch mode; therefore, no images are shown to the user. If you are working with budding yeast or another cell type with a simple round shape, it is advantageous to use the “Convert ROIs to ellipses” option. This approach is also useful for images of nuclei but is not suitable for cells with complex morphologies (e.g., neurons). The “Blind experimenter” option has no effect at this point. If ROIs have already been created, a warning is displayed upon pressing OK, as these will be overwritten if the macro is run, and all changes made previously to the ROIs will be lost. The macro automatically removes objects that are too close to the image edges and are therefore likely to be incomplete.

#### Step 3.2: ROI verification using a Fiji macro

Once the macro is finished, run it again, selecting "Check ROIs" in the process option and using the same options as in the first run. Selecting the “Blind experimenter” option can be used to reduce experimenter bias. Specifically, it randomizes the order in which the images are displayed and hides the filenames and the location of the files from the status message. The channel option specifies which channel is shown to the user. Once OK is pressed, an example of properly made ROIs in yeast cells is displayed to the user as an aid. Then, images of the selected type are displayed one by one to the user together with existing ROIs. These can be removed or adjusted to correspond to the example as closely as possible (the example can be displayed again using the “Help” button in the dialog window). There are options to change the size and position of all ROIs at the same time. To add a new ROI, draw it in the image (the “Make ellipse” tool is preselected) and press ‘t’ on the keyboard to add it to the ROI manager. All changes are automatically saved when OK is pressed in the dialog window. There is an option to return to one of the previous images, or jump ahead, if desired. To increase the visibility of all features, the inverted gray lookup table (LUT) is used to display the images, combined with red outlines of the ROIs. In this step, files can be excluded from the analysis (e.g., if bleached or focused incorrectly) by clicking the “exclude from analysis checkbox”; once OK is pressed, the current image will be closed and moved to the “data-exclude” folder within the current structure. This feature works even when the experimenter is blinded.

*Note that the ROIs defined in this step will be used for the analysis of all available channels in the subsequent steps. If, for example, whole cells are to be analyzed in one channel and nuclei in another, it is recommended to split the images into separate channels and analyze them independently*.

### Step 4: Running the analysis

During the analysis, all images within the specified folder and subfolders are successively opened along with the respective RoiSet (in batch mode, i.e., no images are displayed). Each ROI is then analyzed separately, and the corresponding data is written in a log file that is saved in the form of a csv (comma-separated values) table whenever an image is closed. This enables resumption of an interrupted analysis run. Additional log windows appear, informing the user about the analysis progress (in-cluding rough estimate of the remaining time), listing processed images and images that were not analyzed to absence of defined ROIs.

#### Step 4.1: Running the quantification using a Fiji macro

Once you have checked and adjusted the ROIs, run the “3. Quantify.ijm” macro. In the dialog window that appears, choose the folder from which the analysis is to start (the Results will be saved here). If you only want to process images of a particular yeast strain (cell line) or condition, use the “subset” option – only filenames containing the specified string of characters will be processed. Select which channel(s) you want to analyze (use a comma to specify multiple channels or a dash to define a range; a separate Results table will be created for each channel). Specify the naming scheme you are using for your filenames (information written here will be used as column titles in the Results table). As explained above, the details in the filenames need to be comma-separated to facilitate automatic down-stream filtering and processing. Before running the macro, ensure that the naming is consistent across all the data that are included in the analysis. Specify the format of the experiment code (accession code) you are using and select the proper image type (transversal/tangential). Specify the lower (min) and upper (max) limit for cell area (in μm^2^). Only cells within this range will be included in the analysis. The coefficient of variance option can be used to automatically filter out cells without structure (i.e., probably dead cells). The "Help" button can be used to get information on individual parameters on the fly without needing to consult the protocol. The actual analysis can take a long time, depending on how many images and cells are being analyzed. With big datasets, it is advised to run the analysis overnight (once you have established that there are no issues). Once the analysis is finished a dialog window appears (with a message saying: “Finito!”) with additional information in case of irregularities. The following files are created by the macro:

- “Results of tangential image analysis, channel <CH> (YYYY-MM-DD,HH-MM-SS).csv” or “Results of transversal image analysis, channel <CH> (YYYY-MM-DD,HH-MM-SS).csv” – the "Results table" containing the results of the analysis for the specified channel CH. The numbers in the brackets specify, in the format shown, the date and time when the analysis was concluded.
- <u>“results-temporary_<IMAGE_TYPE>_channel_<CH>.csv”</u> – a temporary file updated every time the analysis of all cells in a single image is concluded. Together with the two files listed below, this file is used to resume an incomplete analysis, e.g., if some images do not have prepared ROIs for analysis, or if the analysis has been interrupted for any reason. This file is converted to the “Results table” file once the analysis of all images is completed.
- <u>“files_without_ROIs.tsv”</u> – lists files from the “data” folder that have no ROIs in the respective “ROIs” folders (and are, therefore, skipped in the analysis). Once these are prepared, the analysis can be resumed (by running it again with the same parameters).
- <u>“processed_files.tsv”</u> – lists the files that have been analyzed. If an incomplete analysis is resumed, these are skipped. *Disclaimer: To count the number of bright fluorescence foci (microdomains), and nowhere else, the analysis uses nonlinear adjustments*.

#### Step 4.2: Examination of the Results table structure

- <u>Header</u> -The Results table contains a header with basic information on the macro run: macro version, date and time when the analysis was started, which channel was analyzed, and a handful of interesting parameters used for the analysis. Each line of the header starts with the pound (#) sign so that it is automatically ignored by the provided R scripts.
- <u>Columns</u> – The first several columns (up to the frame column) contain descriptive parameters, i.e., information about the analyzed sample, derived from the file names. The columns from “mean background” onward contain quantification parameters, i.e., numbers extracted from the microscopy images by the “Quantify” macro. The quantified parameters are described in detail in the “results_table_legend.md” file in our GitHub repository (https://github.com/jakubzahumensky/microscopy_analysis).
- <u>Rows</u> – Each row contains data pertaining to a single cell (ROI)

### Step 5: Extracting numbers, plotting and statistical analysis

The Results table can be further processed in multiple ways. We recommend either taking advantage of the provided R script, which enables fully automated processing, plotting and statistical analysis, or using GraphPad Prism.

#### Using R scripts

Our custom R scripts can be found in our GitHub repository (https://github.com/jakubzahumensky/microscopy_analysis) in the form of a markdown file. They can be used as follows: place the Results table to be processed into a folder together with the provided R markdown file (“microscopy_analysis_data_processing.Rmd”). Open the markdown in R Studio and run the scripts (organized into code chunks with descriptions) – the code can be run all at once using the “Run” button (or by using the “ctrl+alt+R” keyboard shortcut). However, we highly recommend running the individual code chunks separately, at least when the scripts are used for the first time – either by pressing the green “Play” button in the top right corner of each chunk, or by using the “ctrl+shift+enter” keyboard shortcut to run the current code chunk. Detailed descriptions of each analysis step are included in the markdown and guide the user through the whole process step-by-step. A new folder called “analysis” is created with a subfolder for each experiment. The Results table is automatically filtered (grouped) based on all possible unique combinations of descriptive parameters (biological replicates, strains, conditions, etc.). Means and standard deviations (SDs) are calculated and reported in separate tables for each quantified parameter across these groups.

Boxplots showing the median, range, and individual data points for biological replicates are generated for each parameter. The user is prompted to select the independent (x), grouping (factor) and faceting variables from sample description parameters if multiple choices exist. The user can also select quantification parameters for which to plot intricate violin plots, including statistical analyses of differences (Holm-or Bonferroni-corrected multiple *t-tests*).

Independent statistical analysis is performed on data with at least 3 biological replicates (after automatic filtering). Normality (Shapiro‒Wilk) and equal variance (Levene’s test) are tested, followed by multiple unpaired t-tests (using Bonferroni and Holm corrections) and ANOVA with Tukey’s HSD (honestly significant difference) post-hoc test. The results are reported for each quantified parameter in a separate text file.

#### Using GraphPad Prism

Load the Results table by clicking File → New → New Project File → Create Multiple variables and copy-paste the whole Results table here. Click Analyze → Data Processing → Extract and Rearrange, select which parameters you want to extract (we typically use BR_date, strain, condition + the graphed variable, such as cell intensity), select Format of results table (e.g., Grouped for two-way ANOVA) and Data arrangement for plotting and statistical analysis. The exact choices depend on the actual data.

#### Using Microsoft Excel/LibreOffice Calc

When opening the Results file in Excel/Calc, ensure that commas are used as column separators. Once open, select the row with the column headers (exp_code, BR_date, etc.); then, click Data → AutoFilter in Calc or Data → Filter in Excel (a funnel icon in both). Now, clicking on each column title, you can select which lines to display. These can then be moved/copied elsewhere to calculate the mean and standard deviation (SD) within each biological replicate, strain and condition.

*Note: The tables containing means and SDs generated by the R script can be used directly in GraphPad Prism, Systat SigmaPlot (or another scientific graphing software) to create graphs and perform statistical analysis. Before meaningful plots can be generated and statistical analysis performed, certain data (chiefly fluorescence intensity) need to be normalized (see Discussion). As a good guide on how to plot your data, we recommend a recent paper that described what the authors termed SuperPlots. Using these plots, information on individual cells, including their respective biological replicates, is readily visible*^14^.

### Demonstration of data analysis approach versatility

We have previously successfully employed the sample preparation method and Fiji macros described here in multiple studies, validating this approach^7–9^. We have updated the macros after these papers were published to incorporate new features (e.g., resume incomplete analysis, analyze multiple channels in a single run, blind experimenter, additional approaches for the quantification of plasma membrane microdomains) and fix known issues (e.g., macro running commands before the previous one has been completed). In addition, we created R scripts that greatly facilitate downstream processing, graphing and statistical analyses of the data. To demonstrate the universality and robustness of the reported Fiji macros, we include the analysis of a microscopy image of a neuron as an example of a morphologically complex cell (Fig. 8).

**Figure 8.**
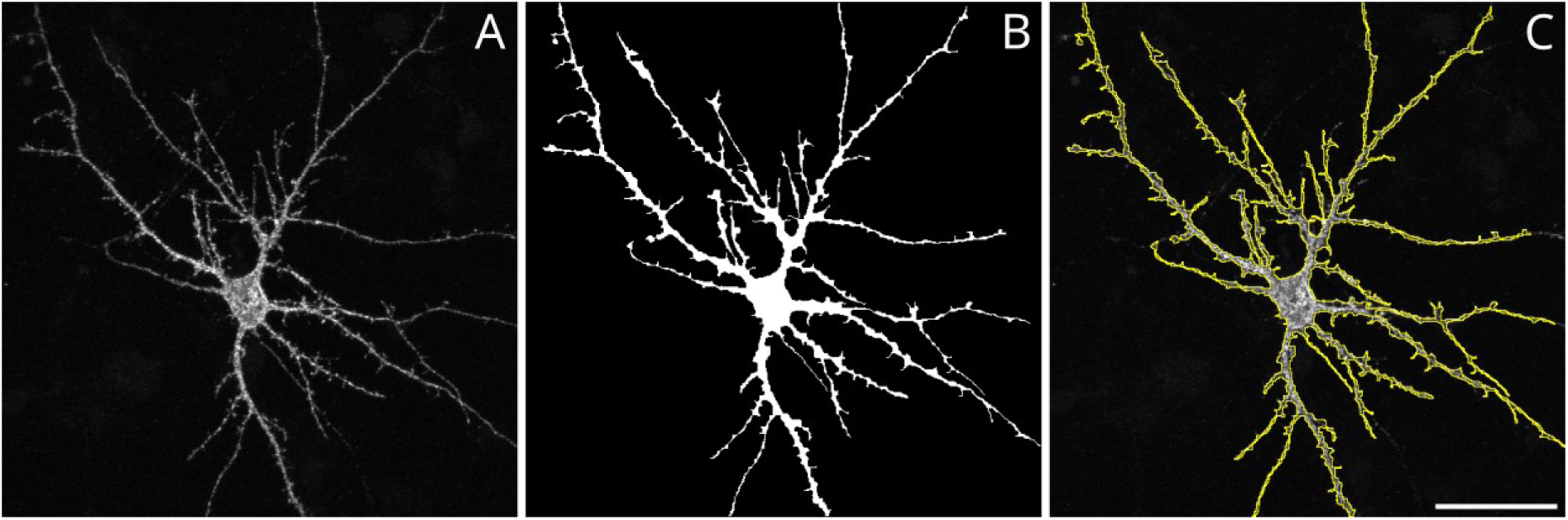
Segmentation of a neuron for protocol validation. A) A z-stack of 21 images was acquired using an Olympus FV10i confocal fluorescence microscope, and maximum intensity projection was calculated in Fiji B) Segmentation mask was created by thresholding image A followed by manual adjustments (Fiji). C) Region of interest (ROI) was created from the mask in B. Primary hippocampal neuronal cultures were prepared from rats at embryonic day 18. The neurons were transfected with a DNA vector containing gene encoding the GFP-GluN3A subunit of N-methyl-D-aspartate receptors on in vitro day 13. Immunofluorescence labeling was performed 24 hours after transfection using primary rabbit anti-GFP and secondary anti-rabbit IgG conjugated Alexa 647 antibodies for surface receptors. The intracellular signal was enhanced using primary mouse anti-GFP and secondary anti-mouse IgG conjugated Alexa 488 antibodies after permeabilization. Scale bar: 50 µm. Image courtesy of Petra Zahumenska. The analysis results are available in our Zenodo repository (https://zenodo.org/records/11517778).

## Discussion

### Key capabilities

We report a simple sample preparation method that enables live cell-walled cells to be gently immobilized in a monolayer, imaged under almost natural conditions and treated directly during a time-lapse experiment. We developed a Fiji-based analysis technique for budding yeast with GFP-tagged proteins that enables batch analysis of large microscopy datasets. Its use is not limited by any imaging platform, microscopy type, labeling or organism, as we demonstrate below. Multiple parameters are reported for each object, including size, fluorescence intensity in the plasma membrane and cytosol and the number of microdomains (i.e., areas of high local intensity). The analysis is largely automated and requires user input only to verify the accuracy of segmentation, with clear rules and an option for blinding this step. Hence, our approach allows for unbiased batch analysis of large datasets, greatly improving the efficiency and accuracy of fluorescence microscopy research. Combined with the provided R scripts, which enable streamlined processing of the data extracted from the microscopy images to report summary data (means and standard deviations), plot graphs and perform statistical analysis, it represents a very powerful tool that can be used wherever microscopy images are acquired and analyzed.

### Comparison of sample preparation methods

The immobilization of cells on microscopy slides is usually achieved by coating these with either poly-L-lysine (PLL) or concanavalin A (ConA). A recent study compared these two options and found that ConA is superior in each analyzed aspect – the required concentration is lower, the immobilization is more effective and more stable, and the protocol takes shorter time. Neither chemical interferes with the cell cycle and budding of yeast, making both suitable for live-cell experiments^15^.

The preparation of ConA-coated microscope slides is straight-forward and takes ∼10 minutes (can be performed at least a day in advance). After cells are applied to the slide, they need to be allowed to settle for ∼ 5 minutes^15^. This creates a delay between cell harvesting and imaging. In addition, ConA is almost completely ineffective for immobilization of cells taken directly from cultures grown in YPD (yeast-peptone-dextrose) and to a lesser degree also SC (synthetic complete) media, i.e., not washed in water or buffer. The reason for this is that YPD components, metabolites and cell debris likely compete for binding to ConA^15^. The need to wash the cells before imaging makes ConA inappropriate for experiments when imaging in the growth media is required and adds to the delay between harvesting and imaging. ConA also does not work after cell wall digestion using Zymolyase and can induce cell aggregation when used at higher concentrations^15^.

Like ConA and PLL use, our approach allows live-cell imaging while not interfering with the cell cycle and budding^7^, with the advantage that the cells can be imaged immediately after the cell suspension is covered with the agarose block. We do, however, recommend that a few minutes are allowed for the cells to settle when a time-lapse experiment is to be performed. Another advantage of our approach is that no slide pretreatment is required before imaging. The agarose is prepared in a 9 cm Petri dish and can be stored (sealed with parafilm) in a fridge for up to a month. Before the experiment, it needs to be moved to room temperature or the incubator to prevent heat stress when applied to the cell suspension. We find the preparation and manipulation of the agarose blocks to be simple and easy to learn. Unlike ConA, the immobilization of cells using agarose blocks is not limited to washed cells and works just as effectively with suspensions made directly from both YPD-and SC-grown cultures. In addition, we were also able to successfully image yeast protoplasts (following 2-hour cell wall digestion with 0.5 mg ml^−1^ Zymolyase 20 T, stabilized with 1 M sorbitol) using our approach^16^.

### Comparison of our image analysis tools with commercial options

Among the advantages of using our custom scripts is that the software required, as well as the codes themselves, are freely accessible to anyone, significantly cutting the cost of analysis. The code is available under the CC BY-NC license, meaning that the users are free to adjust the code and reuse it for other purposes (see “Code availability” for more information). This is facilitated by the extensive use of functions, giving the code a modular character, especially in the case of the Fiji macros. Further Fiji plugins and/or functions can be incorporated into the code, as required. In contrast, in addition for the usually very high initial cost of a license (or a recurring subscription fee), commercial solutions often require the users to pay extra money for expansions of the capabilities of the software, whether it means additional type of analysis, implementation of macros and/or custom code, or the ability to batch-process files. Our workflow has the batch-processing functionality already built in, and this is one of the things that makes it powerful – not only can it process all files within a folder (which is often the limitation of the batch processing options in commercially available solutions), it works recursively. This means that an endless number of folders can be processed in a single run, across unrelated experiments. Since the analysis is automated, it is free of experimenter bias. The only experi-menter-dependent step, i.e., the validation of segmentation, can be blinded on demand, increasing objectiveness of the analysis.

The commercial packages provide a predefined and limited set of parameters that can be analyzed, which invariably include functionalities that the user does not need, while other, desired ones are missing. This can be circumvented by writing custom code and implementing it into the commercial solution, if possible. We developed our workflow based on extensive first-hand experience with microscopy experiments, which is reflected in the choice of parameters that are analyzed and reported. While some parameters may not be directly useful, they are used to calculate other parameters, such as background correction, or intensity ratios. For completeness, we include these parameters in the report (Results table).

The scripts and codes used in this protocol are written in two different languages (Fiji/ImageJ macro language and R). However, they can be used without the need to understand the code or adjust it in any way. Like the commercial solutions, our scripts are meant to work out-of-the box. The running of the scripts is reasonably simple in both Fiji and R Studio. As outlined above, the Fiji macros can be made available in Fiji as menu commands by copying them into a subfolder in the “Fiji.app/scripts” folder, in which case the macros effectively behave as plugins. In R Studio, the code can be run all at once by pressing the “Run” button. However, when used for the first time, we recommend the users to go through the markdown one code chunk at a time, reading the respective descriptions to better understand what the code is doing.

Arguably, the installation process of the software and dependencies required for our workflow is more complicated and takes longer than in the case of commercial solutions. However, the installation needs to be performed only once and the number of users/workstations that can use it is not limited by a license. In contrast to commercial solutions, our workflow does not come with a real graphic user interface (GUI). The only two times the images are displayed to the users are in Cellpose for the determination of segmentation parameters, and in Fiji for verification of segmentation. This is deliberate, as GUIs slow down analyses. Analysis can run in the background and even with a locked screen. The most computing power-demanding part of the analysis is the quantification of plasma membrane microdomains (especially in transversal images). While this parameter is very important for our research, we appreciate that this is not the case in general. Therefore, we incorporated an option to skip this part of the quantification, which significantly speeds up the whole analysis.

Possibly the biggest drawback of our approach lies in the use of multiple Fiji plugins. When these, or Fiji/ImageJ itself, are updated in a fundamental manner, the functionality of the macros may be compromised. This has, indeed, happened to us in the past. However, as we actively use these macros for our work, we are likely to notice such issues promptly and fix them as soon as possible, providing an update. We recommend that users check for updates before performing analysis.

For segmentation of images, we use Cellpose, as we find it easy to use and it gives us consistently accurate results. While it may seem that Cellpose is unnecessarily powerful for our needs, segmenting yeast cells (especially when in close proximity and when budding) is not as simple as it may appear^17^. We have tried various Fiji plugins (Level Sets, StarDist, etc.) but we either found them difficult to use and implement into macros, or they required training of a new model. Invariably, we found the segmentation accuracy inferior to that obtained using Cellpose. Nevertheless, as the segmentation masks are ultimately converted to ROIs before the analysis, the users are free to use their preferred segmentation tool.

### Troubleshooting

#### Sample preparation

For the imaging and subsequent analysis of yeast cells, it is ideal to prepare the microscopy sample in a way that a monolayer is formed. In this case, all cells are localized in a common focal plane and the microscopy images do not contain a lot of empty space. This saves time during both image acquisition and analysis, as a high number of cells is present in each image. In turn, it saves storage space and facilitates inclusion of multiple cells in representative images for publication. Using our approach, where a cell suspension is gently overlayed with an agarose block, yeast cells have natural propensity to organize themselves into monolayers (Fig. 3A, B). However, using a cell suspension that is either too diluted or too dense can lead to issues, resulting in wide gaps between (groups of) cells or multi-layered cell organization (Fig. 3C and D respectively). If the cell suspension density is too low, all cells are in the same focal plane (which is desirable), but a large number of frames needs to be recorded to image a sufficient number of cells (we aim to image at least 200 cells per biological replicate per condition, which enables us to later filter cells for example by cell sizes or to exclude buds from the analysis; dead cells are also not always immediately apparent during imaging and should be excluded from the analysis). This consumes both time and storage space. <u>To circumvent this, w</u>e recommend preparing another microscopy sample after concentrating the cell suspension by a brief and gentle centrifugation (not exceeding 1500 rcf for yeast). The opposite extreme situation takes place when the cell suspension is too dense, leading to formation of multilayers. In this situation, a lot of cells occupy the same x-y coordinates and transversal sections are mixed with tangential sections from another layer in a single focal plane. Imaging such samples is to be avoided. We recommend preparing another sample – the formation of multilayers can be partially avoided by distributing the 1 μL of suspension into 3-4 spots on the cover glass, not applying it whole in a single spot. Alternatively, the cell suspension can be centrifuged and only a part of the pellet gently disturbed by pipetting.

#### Data analysis

There are several issues one can run into when using our analysis macros. The known common pitfalls are listed below and mostly involve incorrect paths to folders. Should any additional issues arise, the readers are welcome to contact the authors.

**Table 1:** Troubleshooting of the most common analysis issues.

| Issue | Possible cause(s) | Solution(s) |
| --- | --- | --- |
| Cellpose does not export Masks properly | The path to the images to be processed contains spaces | Segment all images in a dedicated folder without any spaces anywhere in the path, e.g., “D:\segmentation\” and then move the “masks” folder to the folder of the respective biological replicate |
| Cellpose does not use GPU to calculate the segmentation masks | The used PC is running an AMD-family GPU (under Windows) | There is no solution known to the authors at the moment of writing, the Masks will be computed using the CPU, which will take longer; alternatively, check the Cellpose documentation for new developments on the [topic](https://cellpose.readthedocs.io/en/latest/installation.html#amd-gpu-rocm-installation) (https://cellpose.readthedocs.io/en/latest/installation.html#amd-gpu-rocm-installation). |
|  | Required dependencies are not satisfied. | Follow the instructions in the official Cellpose documentation (https://cellpose.readthedocs.io/en/latest/installation.html#) |
| No data reported in the Results table | The required data structure is not in place. | Check the file structure, consulting Fig. 4. Note that the folders called “transversal”, “tangential” and “data” are absolutely required. |
|  | Cells were not segmented and/or the segmentation masks were not converted into ROIs. In this case, you should get a message informing you that ROIs are missing, and a list of files affected by this. | Check that there is a folder called “ROIs” and that the filenames correspond to those in the respective “data” folder (and have a “-RoiSet.zip” suffix). |
| Currently hypothetical – “file not found” during conversion of Masks to ROIs | After publishing of this protocol, Cellpose developers have changed the naming of the exported segmentation masks. | Check that the filenames end with “_cp_masks.png”. If they do not, replace the value assigned to the “MasksSuffix” variable in line 1 of the “2. ROI_prep.ijm” macro from “_cp_masks.png” to the new suffix (recommended) or rename the Masks files. |

### Note on normalization

Certain values (e.g., the mean fluorescence intensity of a cell) need to be normalized before the statistical significance of the observed changes between strains/cell lines and/or conditions/treatments can be tested. The usual practice is to simply divide values belonging to a single biological replicate set by the value of the control sample. While useful, it hides the variability of the control sample itself and lowers the power of the statistical tests employed. To circumvent this, we suggest the following normalization procedure and provide an example (Table 2; assuming the same set of conditions is used in each biological replicate):

1. Calculate the mean of all values within a biological replicate set.
2. Divide each value within the biological replicate set by this value.
3. Calculate the mean value of the control samples across replicates.
4. Divide all scaled values across replicates by the mean of the controls.

**Table 2.** Normalization example. The raw data are first scaled within their respective biological replicate groups and then normalized using the mean value of the control samples.

| condition | raw data |  |  | scaled |  |  | normalized to control |  |  |
| --- | --- | --- | --- | --- | --- | --- | --- | --- | --- |
|  | replicate 1 | replicate 2 | replicate 3 | replicate 1 | replicate 2 | replicate 3 | replicate 1 | replicate 2 | replicate 3 |
| control | 20 | 25 | 50 | 0.83 | 0.80 | 0.86 | 1.00 | 0.96 | 1.04 |
| treatment 1 | 25 | 38 | 64 | 1.04 | 1.22 | 1.10 | 1.25 | 1.46 | 1.33 |
| treatment 2 | 21 | 22 | 48 | 0.88 | 0.70 | 0.83 | 1.05 | 0.85 | 0.99 |
| treatment 3 | 30 | 40 | 70 | 1.25 | 1.28 | 1.21 | 1.50 | 1.54 | 1.45 |

## Author contributions

**Jakub Zahumensky** – Conceptualization, Data curation, Formal analysis, Investigation, Methodology, Project administration, Software, Validation, Visualization, Writing -original draft, Writing - review & editing. **Jan Malinsky** – Conceptualization, Funding acquisition, Methodology, Resources, Supervision, Validation, Writing - review & editing

## Acknowledgments

Microscopy was performed at the Microscopy Service Centre of the Institute of Experimental Medicine CAS supported by the MEYS CR (LM2023050 Czech-Bioimaging). The Fiji macros provided here were originally written for our previous publications^7–9^. We further acknowledge our colleagues Petra Zahumenska, MSc for providing us with a sample neuron image, and Dr Petra Vesela for helping with the preparation of images depicting detailed sample preparation and critical input on the manuscript.

## Data and code availability

Code described in this article and sample data have been deposited to GitHub: https://github.com/jakubzahumensky/microscopy_analysis and Zenodo (https://zenodo.org/records/11517778), respectively. The code is available under the CC BY-NC license, i.e., it is open source, and users are free to use and modify it according to their needs, as long as it is not used for commercial purposes. We kindly ask users that if they use our code in either its original or modified form, that a citation to this Protocol be included in their publications. The code is under active maintenance, and we recommend users to check for updates and new features before per-forming the analysis.

## References

1. Gournas, C. et al. Transition of yeast Can1 transporter to the inward-facing state unveils an α-arrestin target sequence promoting its ubiquitylation and endocytosis. Mol. Biol. Cell 28, 2819– 2832 (2017).

2. Chen, C. H. & Pan, C. L. Live-cell imaging of PVD dendritic growth cone in post-embryonic C. elegans. STAR Protoc. 2, 100402 (2021).

3. Grousl, T., Opekarová, M., Stradalova, V., Hasek, J. & Malinsky, J. Evolutionarily conserved 5’-3’ exoribonuclease Xrn1 accumulates at plasma membrane-associated eisosomes in post-diauxic yeast. PLoS One 10, e0122770 (2015).

4. Vaškovičová, K. et al. mRNA decay is regulated via sequestration of the conserved 5′-3′ exoribonuclease Xrn1 at eisosome in yeast. Eur. J. Cell Biol. 96, 591–599 (2017).

5. Grossmann, G., Opekarová, M., Malinsky, J., Weig-Meckl, I. & Tanner, W. Membrane potential governs lateral segregation of plasma membrane proteins and lipids in yeast. EMBO J. 26, 1–8 (2007).

6. Stringer, C., Wang, T., Michaelos, M. & Pachitariu, M. Cellpose: a generalist algorithm for cellular segmentation. Nat. Methods 18, 100–106 (2021).

7. Zahumenský, J. et al. Microdomain Protein Nce102 Is a Local Sensor of Plasma Membrane Sphingolipid Balance. Microbiol. Spectr. 10, e0248922 (2022).

8. Balazova, M. et al. Two Different Phospholipases C, Isc1 and Pgc1, Cooperate To Regulate Mitochondrial Function. Microbiol. Spectr. 10, e0248922 (2022).

9. Vesela, P., Zahumensky, J. & Malinsky, J. Lsp1 partially substitutes for Pil1 function in eisosome assembly under stress conditions. J. Cell Sci. 136, jcs260554 (2023).

10. Zahumenský, J. & Malínský, J. Live cell microscopy sample preparation (yeast culture). protocols.io 1–7 (2024) doi:10.17504/protocols.io.8epv5r23dg1b/v1.

11. Brown, C. M. Fluorescence microscopy – avoiding the pitfalls. J. Cell Sci. 120, 3488–3488 (2007).

12. Sage, D. & Unser, M. Easy Java programming for teaching image-processing. in Proceedings 2001 International Conference on Image Processing (Cat. No.01CH37205) vol. 2 298–301 (IEEE, 2001).

13. Stirling, D. R. et al. CellProfiler 4: improvements in speed, utility and usability. BMC Bioinformatics 22, 1–11 (2021).

14. Lord, S. J., Velle, K. B., Dyche Mullins, R. & Fritz-Laylin, L. K. SuperPlots: Communicating reproducibility and variability in cell biology. J. Cell Biol. 219, (2020).

15. Caloca, B. et al. Comparison of concanavalin A and poly-<scp>l</scp> -lysine as cell adhesives for routine yeast microscopy applications. Yeast 39, 312–322 (2022).

16. Malinska, K., Malinsky, J., Opekarova, M. & Tanner, W. Distribution of Can1p into stable domains reflects lateral protein segregation within the plasma membrane of living S. cerevisiae cells. J. Cell Sci. 117, 6031–6041 (2004).

17. Dietler, N. et al. A convolutional neural network segments yeast microscopy images with high accuracy. Nat. Commun. 11, 1–8 (2020).

